# Ponatinib shows potent antitumor activity in small cell carcinoma of the ovary hypercalcemic type (SCCOHT) through multi-kinase inhibition

**DOI:** 10.1101/159905

**Authors:** Jessica D. Lang, William P. D. Hendricks, Holly Yin, Jeffrey Kiefer, Pilar Ramos, Ritin Sharma, Patrick Pirrotte, Elizabeth A. Raupach, Chris Sereduk, Nanyun Tang, Winnie Liang, Megan Washington, Salvatore J. Facista, Victoria L. Zismann, Emily M. Cousins, Michael B. Major, Yemin Wang, Anthony N. Karnezis, Krystal A. Orlando, Aleksandar Sekulic, Ralf Hass, Barbara Vanderhyden, Kesavannair Praveen, Bernard E. Weissman, David G. Huntsman, Jeffrey M. Trent

## Abstract

Purpose: Subunits of the SWI/SNF chromatin-remodeling complex are tumor suppressors inactivated in ∼20% of all cancers. Yet, few targeted treatments for SWI/SNF-mutant cancers exist. Small cell carcinoma of the ovary, hypercalcemic type (SCCOHT) is a rare, aggressive ovarian cancer in young women that is universally driven by loss of the SWI/SNF ATPase subunits, SMARCA4 and SMARCA2. Given poor two-year survival rates for these women, a great need exists for effective targeted therapies.

Experimental Design: To identify underlying therapeutic vulnerabilities in SCCOHT, we conducted high-throughput siRNA and drug screens. Complementary proteomics approaches comprehensively profiled kinases inhibited by ponatinib. Ponatinib was tested for efficacy in two PDX models and one cell line xenograft model of SCCOHT.

Results: FGFRs and PDGFRs were overlapping hits between screens and the receptor tyrosine kinase (RTK) family was enriched in the siRNA screen hits. Evaluation of eleven RTK inhibitors in three SCCOHT cell lines identified ponatinib, an inhibitor of multiple RTKs, as the most effective clinically approved agent. Proteomics approaches confirmed inhibition of known targets of ponatinib and more than 20 non-canonical ponatinib targets. Ponatinib also delayed tumor doubling time 4-fold in SCCOHT-1 xenografts and reducing final tumor volumes in two SCCOHT patient-derived xenograft (PDX) models by 58.6% and 42.5%.

Conclusion: Ponatinib is an effective agent for SCCOHT in both *in vitro* and *in vivo* preclinical models through its inhibition of multiple kinases. Clinical investigation of this FDA-approved oncology drug in SCCOHT is warranted.

**Additional Information:** This work was supported by research funds from the Canadian Cancer Society Research Institute 34 (#703458, D.G.H.), the National Institutes of Health (R01 CA195670-01, B.E.W., D.G.H., and 35 J.M.T., and T32 HL007106-39 to E.M.C), the Terry Fox Research Institute Initiative New Frontiers Program in Cancer (#1021, D.G.H.), the British Columbia Cancer Foundation (D.G.H.), the VGH & UBC Foundation (D.G.H.), the Anne Rita Monahan Foundation (P.R.), the Marsha Rivkin Center for Ovarian Cancer Research (J.M.T.), the Ovarian Cancer Alliance of Arizona (J.M.T.), the Small Cell Ovarian Cancer Foundation (P.R., J.D.L., B.V., and J.M.T.), and philanthropic support to the TGen Foundation (J.M.T.).

COI disclosure statement: The authors declare no potential conflicts of interest.

## Statement of Translational Relevance

Pathogenic mutations in SWI/SNF chromatin-remodeling complex members occur in approximately 20% of cancers, but no targeted therapies that exploit a tumor’s dependence on SWI/SNF dysfunction have yet shown clinical impact. Small cell carcinoma of the ovary, hypercalcemic type (SCCOHT) is a rare and aggressive form of ovarian cancer affecting young women. It is characterized by mutational inactivation of SMARCA4 and epigenetic silencing of SMARCA2, the mutually exclusive ATPases of the SWI/SNF complex. Here, we demonstrate potency of the FDA-approved oncology drug, ponatinib, a receptor tyrosine kinase (RTK) inhibitor whose targets include PDGFRs, FGFRs and EphAs, in SCCOHT cell and animal models. This work suggests that ponatinib exploits SCCOHT’s dependence on RTK signaling in the context of SWI/SNF dysregulation and that ponatinib may be effective in SCCOHT treatment. Preclinical identification of an effective, approved oncology drug holds promise for rapidly improving outcomes for these young patients and warrants clinical investigation.

## Introduction

Small cell carcinoma of the ovary, hypercalcemic type (SCCOHT), a rare and aggressive form of ovarian cancer, is diagnosed in women at a median age of 24 years (range: 14 months to 47 years) (1). Meta-analysis of 257 clinically annotated SCCOHT cases has shown a dismal two-year survival rate less than 35% and the most effective treatment regimen based on this assessment is surgery followed by aggressive high dose chemotherapy, radiation, and stem cell rescue (1,2). The poor response rates and extreme toxicity of this regimen necessitate identification of effective, targeted treatments for these young patients. SCCOHTs are characterized by inactivating germline and somatic mutations in the tumor suppressor *SMARCA4* (also known as BRG1) resulting in concomitant protein loss in nearly all cases (3-8). These *SMARCA4* alterations occur amidst otherwise diploid SCCOHT genomes and very rare secondary mutations in other cancer genes (4,8). SMARCA4 is one of two mutually exclusive ATPase subunits of the SWI/SNF chromatin-remodeling complex that plays a central role in regulation of transcriptional programs associated with differentiation. The alternative SWI/SNF ATPase, SMARCA2 (also known as BRM), is also absent in SCCOHT due to epigenetic silencing (3,7). Thus, these tumors are driven by a unique genotype that fuels broad transcriptional dysregulation through SWI/SNF dysfunction.

Several other tumor types are also universally characterized by inactivation of SWI/SNF complex members including thoracic sarcomas bearing *SMARCA4* mutation and SMARCA2 loss (9), rhabdoid tumors which are universally characterized by *SMARCB1* (also known as SNF5) mutations alongside SMARCA2 silencing and expression loss in 70% of cases (10,11), and renal medullary cancers also characterized by SMARCB1 loss (12,13). Other cancers with a significant proportion of SWI/SNF mutations include ovarian clear cell carcinomas and endometrioid carcinomas (∼50% and ∼30% with ARID1A loss, respectively) (14,15) and non-small cell lung cancers (∼10% of primary tumors with dual SMARCA4 and SMARCA2 loss) (16,17). Overall, an estimated 20% of human cancers bear potentially oncogenic mutations in one or more SWI/SNF complex subunits (18,19). Thus, identification of therapeutic vulnerabilities in SWI/SNF-mutant cancers with relatively simple genomes such as SCCOHT may hold broader relevance for more diverse cancers.

Preclinical studies to date have suggested that several experimental agents such as foretinib (c-Met inhibitor), epothilone B (tubulin inhibitor), or oncolytic viruses may be effective in SCCOHT (20-22). We and others have also shown that investigational epigenetic agents such as bromodomain and EZH2 inhibitors may hold promise for treatment of these cancers (23-25). Yet, despite the prevalence of pathogenic SWI/SNF mutations in cancer, no approved targeted cancer drugs have yet shown activity in the setting of loss of these tumor suppressors. In order to identify novel therapeutic vulnerabilities conferred by SWI/SNF dysfunction in SCCOHT with a focus on identification of targeted FDA-approved oncology drugs, we performed high-throughput (HT) siRNA and drug screens in SCCOHT cell lines. We thereby identified enrichment for dependence on receptor tyrosine kinase (RTK) signaling both via an abundance of RTK hits in the siRNA screen as well as discovery of SCCOHT hypersensitivity to the RTK inhibitor (RTKi) PD-161570 in the chemical screen. Subsequent evaluation of a panel of FGFR/PDGFR-selective RTKis highlighted ponatinib as the most potent tested agent in SCCOHT cell lines. This observation aligns with prior data in SWI/SNF-mutant rhabdoid tumor models in which expression of the ponatinib target FGFR was shown to be associated with SMARCB1 and SMARCA4 loss, conveying sensitivity to RTKis including ponatinib (26,27). Further, activation of downstream AKT was shown to be elevated in SMARCB1-deficient tumor models (28,29) and dual inhibition of PDGFRα and FGFR1 in MRT cell lines was more effective than specific inhibitors (30). Here, we show that ponatinib’s effects in SCCOHT are also mediated by dependence on signaling of multiple RTKs. Finally, we show the ability of ponatinib to significantly delay tumor progression in a cell line xenograft model of SCCOHT and markedly reduce tumor growth of patient-derived xenograft (PDX) models of SCCOHT, thereby prioritizing ponatinib for a clinical trial in SCCOHT patients.

## Materials and Methods

### Cell lines

BIN67, SCCOHT-1, and COV434 (SCCOHT), G401 and G402 (MRT) and H522 (lung adenocarcinoma) cells were maintained in RPMI 1640 (Thermo Fisher Scientific, Waltham, MA, USA) supplemented with 10% Fetal Bovine Serum (FBS; Thermo Fisher Scientific) and 1% Penicillin/Streptomycin (Thermo Fisher Scientific). COV434 cells, previously identified as derived from a juvenile granulosa cell tumor have now been re-categorized as SCCOHT based on SMARCA4 mutation and lack of SMARCA2 expression (24). A427 (lung adenocarcinoma) and HepG2 (hepatocellular carcinoma) cells were maintained in EMEM (Thermo Fisher Scientific) supplemented with 10% FBS and 1% Penicillin/Streptomycin. COV434 (SCCOHT) cells were maintained in DMEM (Thermo Fisher Scientific) supplemented with 10% FBS and 1% Penicillin/Streptomycin. SVOG3e (SV40-transformed ovarian cells) cells were maintained in DMEM/F12 (Thermo Fisher Scientific) supplemented with 10% FBS and 1% Penicillin/Streptomycin. All cells were maintained at 37°C in a humidified incubator containing 5% CO_2_. All cell lines were routinely monitored for mycoplasma testing and STR profiled for cell line verification.

### Western blotting

Whole-cell extracts from cell lines were prepared using RIPA buffer (Santa Cruz Biotechnology, Dallas, TX, USA) with protease and phosphatase inhibitors using standard protocols. Thirty µg protein was loaded per well on NuPage 4-12% Bis-Tris gels and subsequently transferred to PVDF membranes. Blots were pre-blocked in 5% non-fat dry milk in TBST or BSA in TBST for 1 hour, and probed using primary antibody overnight. Blots were incubated with secondary antibody (anti-rabbit IgG-HRP, Cell Signaling Technology, Danvers, MA, USA; anti-mouse IgG-HRP, Santa Cruz Biotechnology) at 1:5000 for 2 hours and developed using Pierce ECL Western Blotting Substrate or SuperSignal West Femto Substrate (Thermo Fisher Scientific). Primary antibodies (Cell Signaling Technology): phospho-p38, total p38, phospho-Akt T308, phospho-Akt S473, total Akt, GAPDH.

### Phospho-RTK profiling

BIN67 and SCCOHT-1 cells were treated with ponatinib at IC_30_, IC_50_, and 1µM for 1 hour. Phospho-RTK profiling was performed using the Proteome Profiler Human Phospho-RTK Array Kit (R&D Systems, Minneapolis, MN, USA) according to manufacturer’s recommendations. Briefly, cells were lysed and the BCA assay was performed to quantify protein concentration. Three hundred μg of lysate was incubated on each blot overnight, and bound phosphorylated protein was detected using HRP-conjugated anti-phospho-tyrosine antibody. Arrays were developed using Pierce ECL Western Blotting Substrate (Thermo Fisher Scientific), and images were quantified using ImageJ software (NIH) using built-in gel quantification and background subtraction tools. Duplicate spots were averaged for each RTK, and ponatinib-treated samples were normalized to vehicle-treated samples.

### Animal studies

All procedures were carried out under the institutional guidelines of TGen Drug Development’s Institutional Animal Care and Use Committee or the Animal Care Committee of the University of British Columbia (SCCOHT-1 model, A14-0290). For SCCOHT-1 xenograft experiments, 1×10^7^ cells in 50% Matrigel / 50% Media (Corning) in a final volume of 200 µl were subcutaneously inoculated into the backs of NRG (NOD.Cg-Rag1^tm1Mom^ Il2rg^tm1Wjl^/SzJ) mice. For PDX experiments, histologically confirmed SMARCA4-mutant SCCOHT tumors PDX-465 and PDX-040 were acquired from Molecular Response (MRL, San Diego, CA, USA) and serially passaged in mice (3). Tumor suspension in 50% Matrigel / 50% Media in a final volume of 100 μL were subcutaneously inoculated into the backs of NOG (NOD.Cg-*Prkdc*^*scid*^*Il2rg^tmlSug^*/JicTac) mice. Mice were randomized to treatment arms (n=8) once the average tumor volume reached 100-400mm^3^ (SCCOHT-1 model) or 75-125 mm^3^ (PDX models). Ponatinib (Selleckchem, Houston, TX, USA or Activebiochem (SCCOHT-1)) was formulated in 25mM citrate buffer (pH 2.75). Vehicle or ponatinib (15 or 30 mg/kg for SCCOHT-1 or PDX models, respectively) was administered by oral gavage daily for 30 days or until humane endpoint (tumor reaches 1000mm^3^). Tumor size and body weight were measured twice weekly until endpoint. Tumors were excised upon necropsy and either frozen or formalin-fixed/paraffin-embedded.

Chemical library, siRNA screens, ABPP, and RNA-Seq methods are described in Supplemental Materials and Methods.

## Results

### *High-throughput* in vitro *siRNA and drug screens prioritize ponatinib as an FDA-approved oncology drug with potent activity in SCCOHT cell lines*

In an effort to identify SCCOHT’s therapeutic vulnerabilities, HT siRNA libraries were used to test necessity of clinically actionable genes or growth of the SCCOHT cell line BIN67. We transfected siRNAs targeting over 7,000 genes in both Druggable Genome (DGv3) and also custom kinome (vKINv4) libraries. Hits were determined based on measurement of cell viability reduction using CellTiter Glo measurements analyzed via a combination of redundant siRNA activity score (RSA) p-value ranking with ranking by mean signal intensity and utilizing a p-value cutoff of 0.05 (see Supplemental Materials and Methods for further detail). Of 246 genes from the DGv3 library and 81 genes from the vKINv4 library identified as hits based on these criteria, 109 were also validated through independent confirmation that two of four siRNAs inhibited viability by >50% (See Supplemental Materials and Methods and Supplemental Table 1). The ReactomeFI Cytoscape plugin was then used to identify pathways enriched in these validated hits (Table 1) (31). Top enriched gene sets were dominated by receptor tyrosine kinase (RTK) signaling, including FGFRs, PDGFRs, and EGFRs alongside the AKT and MAPK signaling cascades downstream of these RTKs. In the protein interaction network generated from gene hits using ReactomeFI, RTK signaling similarly emerged as the central connected network, as demonstrated by the PDGFR signaling pathway (Figure 1A). All kinase hits from the two siRNA screens were also plotted on a kinome tree to identify key kinase families on which SCCOHT cells are reliant for growth (Supplemental Figure 1). Tyrosine kinases, particularly RTKs, were most strongly identified amongst the kinase hits. A second pathway analysis method, ClueGO, was used to visualize the biological concepts represented in the top hits (Supplemental Figure 2). This method also identified interaction between RTKs in top hits through functions in cancer signaling, regulation of actin cytoskeleton, and cell adhesion. Together, these data suggest a broad dependency of SCCOHT cells on RTK signaling networks.

**Table 1.**
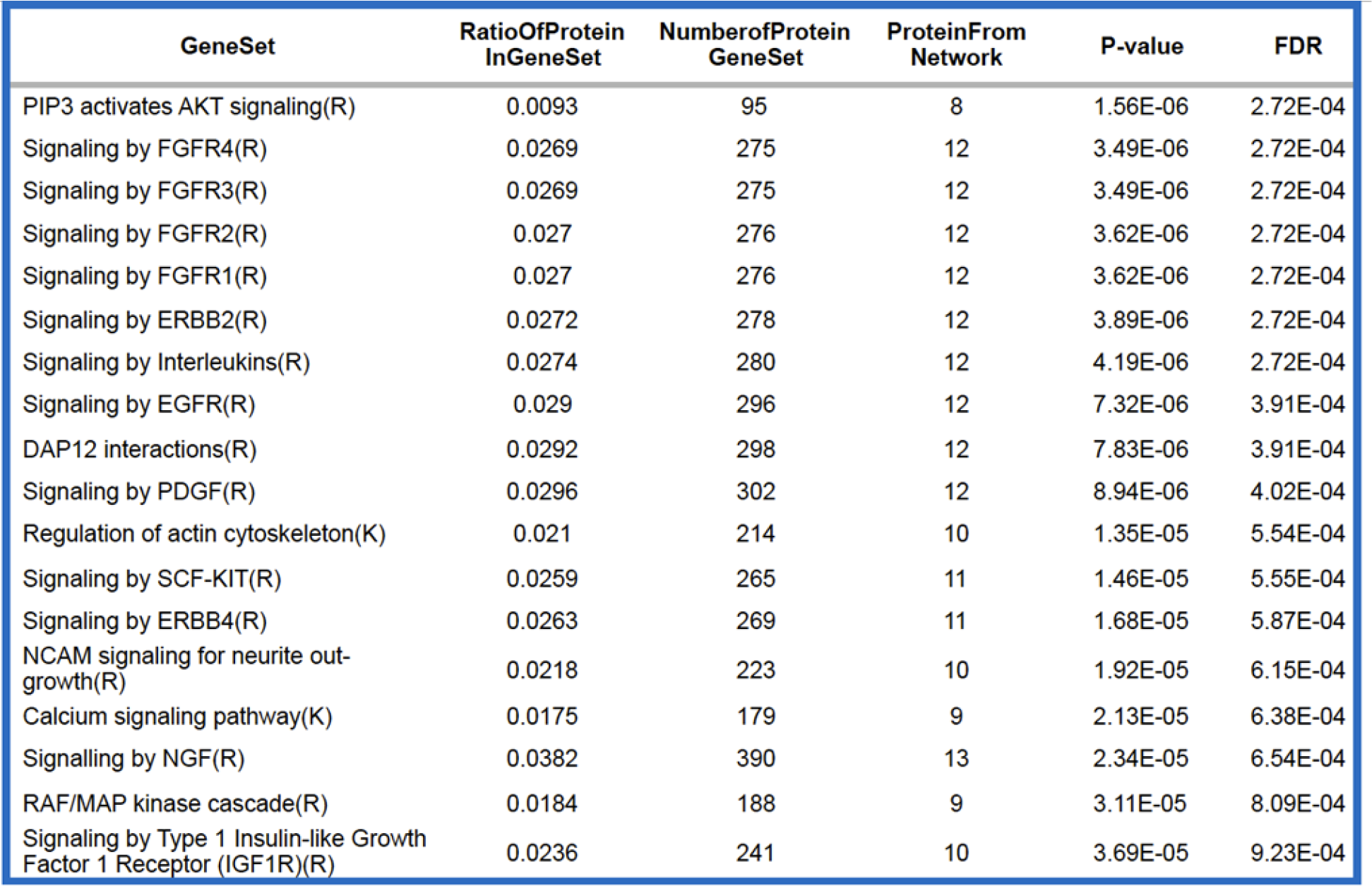
Top pathways enriched in siRNA screen gene hits using ReactomeFI (Cytoscape).

| GeneSet | RatioOfProtein<br>InGeneSet | NumberOfProtein<br>GeneSet | ProteinFrom<br>Network | P-value | FDR |
| --- | --- | --- | --- | --- | --- |
| PIP3 activates AKT signaling(R) | 0.0093 | 95 | 8 | 1.56E-06 | 2.72E-04 |
| Signaling by FGFR4(R) | 0.0269 | 275 | 12 | 3.49E-06 | 2.72E-04 |
| Signaling by FGFR3(R) | 0.0269 | 275 | 12 | 3.49E-06 | 2.72E-04 |
| Signaling by FGFR2(R) | 0.027 | 276 | 12 | 3.62E-06 | 2.72E-04 |
| Signaling by FGFR1(R) | 0.027 | 276 | 12 | 3.62E-06 | 2.72E-04 |
| Signaling by ERBB2(R) | 0.0272 | 278 | 12 | 3.89E-06 | 2.72E-04 |
| Signaling by Interleukins(R) | 0.0274 | 280 | 12 | 4.19E-06 | 2.72E-04 |
| Signaling by EGFR(R) | 0.029 | 296 | 12 | 7.32E-06 | 3.91E-04 |
| DAP12 interactions(R) | 0.0292 | 298 | 12 | 7.83E-06 | 3.91E-04 |
| Signaling by PDGF(R) | 0.0296 | 302 | 12 | 8.94E-06 | 4.02E-04 |
| Regulation of actin cytoskeleton(K) | 0.021 | 214 | 10 | 1.35E-05 | 5.54E-04 |
| Signaling by SCF-KIT(R) | 0.0259 | 265 | 11 | 1.46E-05 | 5.55E-04 |
| Signaling by ERBB4(R) | 0.0263 | 269 | 11 | 1.68E-05 | 5.87E-04 |
| NCAM signaling for neurite out-<br>growth(R) | 0.0218 | 223 | 10 | 1.92E-05 | 6.15E-04 |
| Calcium signaling pathway(K) | 0.0175 | 179 | 9 | 2.13E-05 | 6.38E-04 |
| Signalling by NGF(R) | 0.0382 | 390 | 13 | 2.34E-05 | 6.54E-04 |
| RAF/MAP kinase cascade(R) | 0.0184 | 188 | 9 | 3.11E-05 | 8.09E-04 |
| Signaling by Type 1 Insulin-like Growth<br>Factor 1 Receptor (IGF1R)(R) | 0.0236 | 241 | 10 | 3.69E-05 | 9.23E-04 |

**Figure 1.**
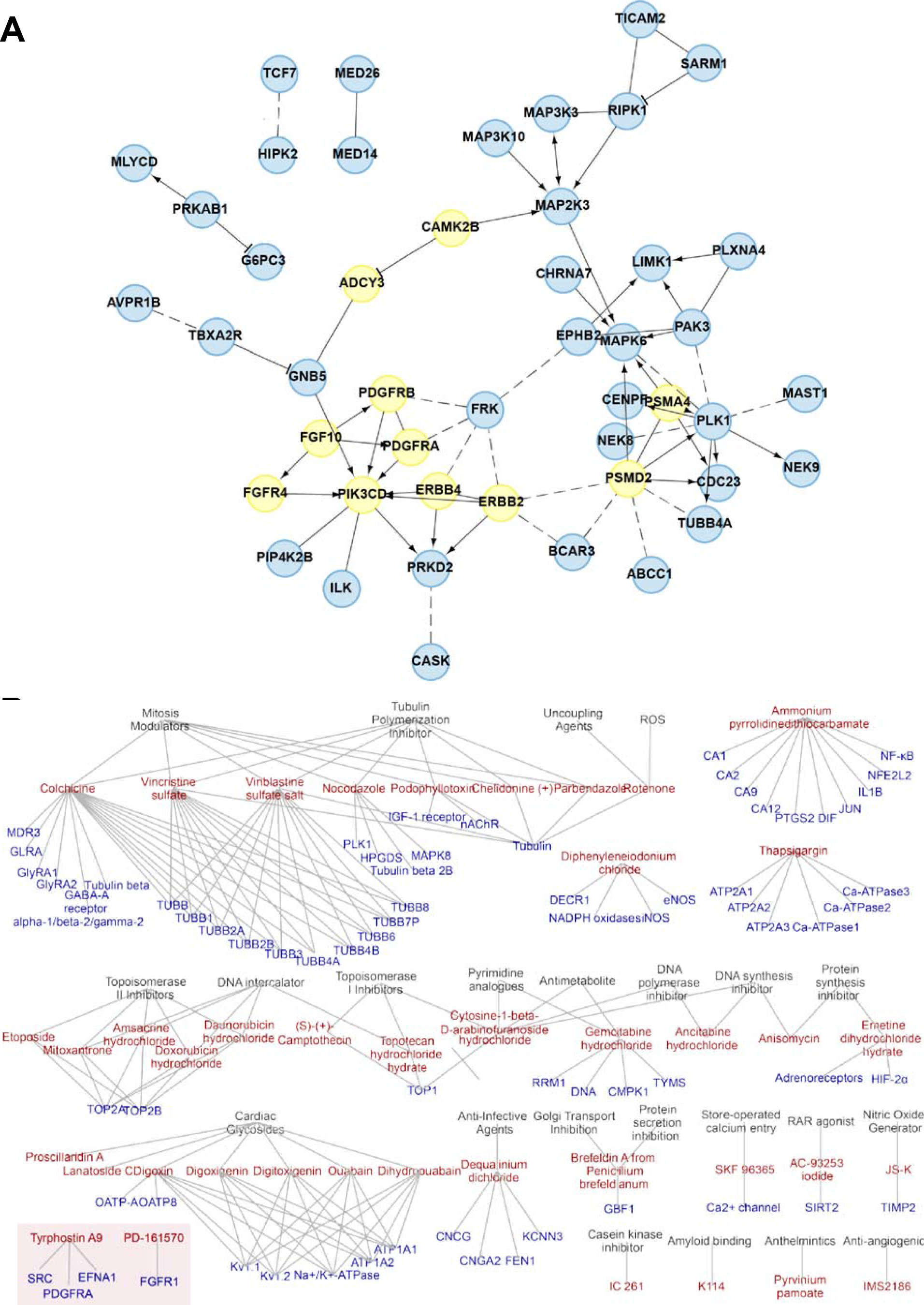
High-throughput screen network analysis. (A) Protein interaction network constructed for RNAi validated screen hits using ReactomeFI network resources. Edges connected to nodes represent evidence for an association between two nodes, which represent the validated RNAi gene hits. Nodes with no edges are not pictured here (full validated hit list in Supplemental Table 3). (B) Chemical hit entity association network for verified chemical hits and their targets and associated biological concepts in SCCOHT cell line, BIN67. RTK associated hits are highlighted in box.

In an effort to identify novel candidate drugs for targeted treatment of SCCOHT in an unbiased manner, high-throughput (HT) drug screens were performed in BIN67 cells using two libraries of ∼2,300 small molecules. All of these compounds are pharmacologically active with well-annotated drug targets from a wide breadth of signaling pathways, and over half of these are approved for at least one clinical indication. Given that SCCOHT’s cell of origin remains unknown and, further, that isogenic model systems restoring SMARCA4 expression in SCCOHT result in senescence, we did not utilize a normal or isogenic comparator to identify selective hits. Instead, we chose to identify compounds active in BIN67 SCCOHT cells and to filter out broadly cytotoxic drugs that also showed effect in HepG2 cells. Hits were then defined as those agents that reduced viability of BIN67 cells by >50% relative to HepG2 according to CellTiter Glo measurements at 72 hours post-treatment (see Supplemental Materials and Methods for further detail). From these screens, 64 compounds reduced growth of BIN67 but not HepG2 cells (Supplemental Table 2). Of these hits, 51 were next validated in 12-point drug dose-response (DDR) format in the SCCOHT cell lines BIN67, SCCOHT-1, and COV434 as well as a SMARCA4-wild-type human granulosa cell line SVOG3e and HepG2 cells. Of the 51 compounds tested, 42 were confirmed to have IC_50_ values >10-fold lower in BIN67 cells than HepG2 cells (Supplemental Table 3). Generally IC_50_ values from BIN67 cells were similar to those in SCCOHT-1 and COV434, as well as for SVOG3e. The 42 validated drug hits were annotated based on their reported targets in preclinical literature and vendor database annotations. This information was incorporated into a chemical-target/entity bipartite network and visualized using Cytoscape (Figure 1B). Network cliques with the highest number of connected compounds were associated with microtubule targeting or DNA damage induction. These network cliques are representative of current standard treatments for SCCOHT, which include vinblastine, doxorubicin, and etoposide (2). While the original Prestwick chemical library did not contain any annotated RTKi, the LOPAC library included 27 RTKi, of which nine were EGFR inhibitors, two were IGF-1R inhibitors, two were FGFR inhibitors, one was a PDGFR inhibitor, and the remaining targeted miscellaneous or unannotated RTKs. Two of these RTKi were identified as hits in the validation set: PD-161570 and Tyrphostin A9. While these agents demonstrate some selectivity for FGFR1 and PDGFRs, respectively, they are not selective for these targets alone, and are known to broadly target multiple RTKs.

To follow up on evidence of RTK dependence in the siRNA and chemical screens, we assessed sensitivity of BIN67 and SCCOHT-1 to 11 RTKi. We selected a subset of RTKi, particularly those targeting FGFRs and PDGFRs. Drug Dose Response (DDR) assays were performed (Figure 2A and B) in an expanded panel of cell lines with known defects in SWI/SNF complex members including SMARCB1- and SMARCA2-deficient MRTs (G401 and G402) and SMARCA4 and SMARCA2-deficient lung cancer cell lines (A427 and H522). Overall, these lines were most sensitive to the broad-spectrum RTKi PD166285 and ponatinib. Ponatinib was further prioritized based on existing approval for clinical use in chronic myeloid leukemia and acute lymphoblastic leukemia. Of known ponatinib targets including *FLT3, KIT*, FGFRs, PDGFRs, *RET*, and *ABL* (32-34), only *FGFR4, PDGFRA*, and *PDGFRB* were validated siRNA screen hits in BIN67 cells (Figure 2C).

**Figure 2.**
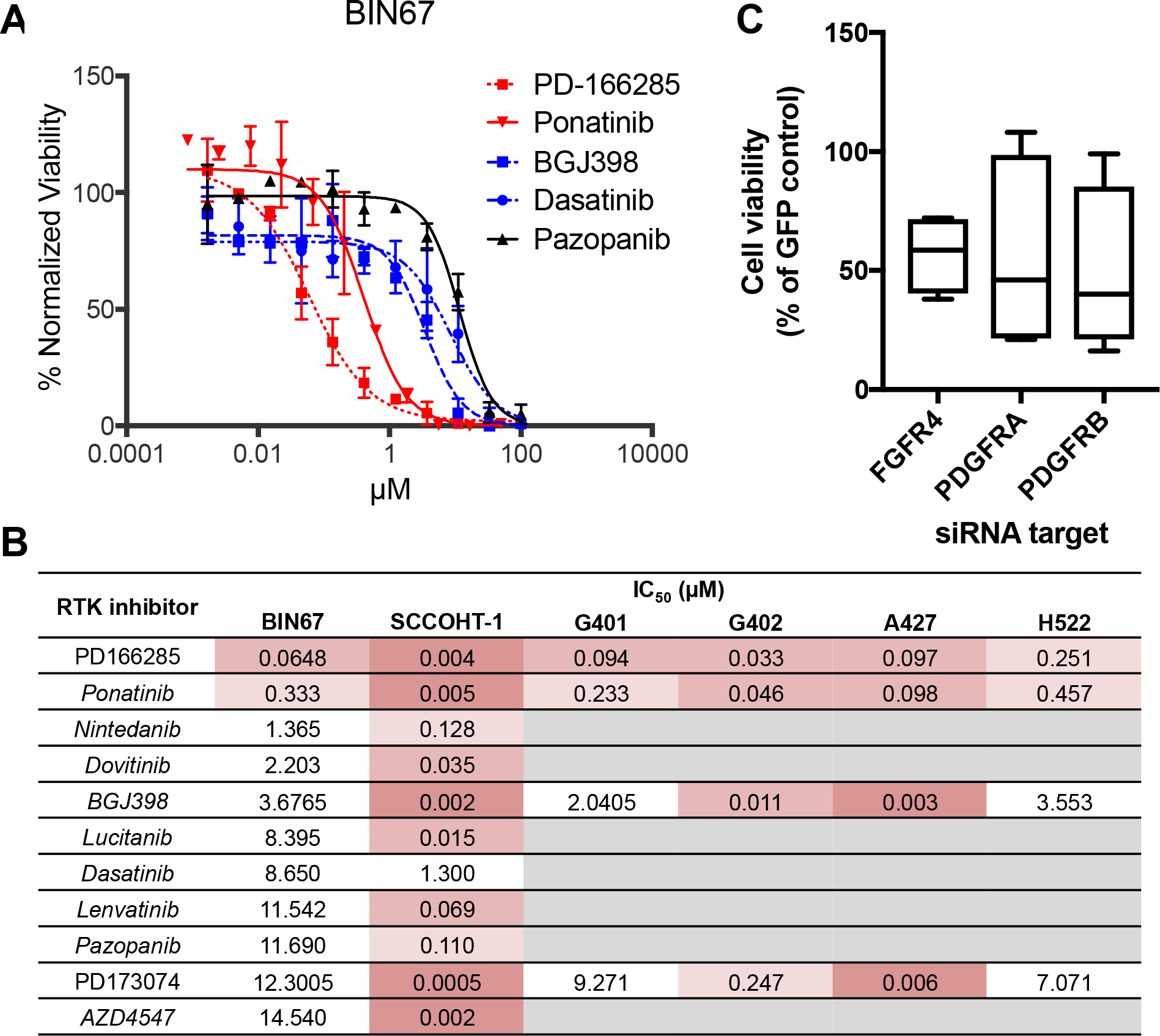
Sensitivity of SCCOHT and other SWI/SNF mutant cell lines to RTK inhibition. (A) DDR curves for various RTKi in BIN67 cells. Cell viability was assayed after hours treatment. (B) Summary of IC_50_ values for RTKi tested in SCCOHT cell lines, along with selected agents tested in SWI/SNF-mutant MRT cell lines (G401 and G402) and lung cancer cell lines (A427 and H522). Conditions not tested are in gray. (C) Cell viability of BIN67 cells following knockdown of ponatinib target genes. Known ponatinib target genes that were initial hits in the siRNA screen (FGFR4, PDGFRA, and PDGFRB) were validated in a secondary siRNA screen. Data shown is mean cell viability of 3 independent replicates of 4 distinct siRNAs relative to internal GFP-targeting siRNA control. All three target genes were considered validated hits based on criteria described in methods.

### Known RTK targets of ponatinib are expressed in SCCOHT tumors and downstream signaling is inhibited in response to ponatinib treatment

Because ponatinib is known to broadly target several RTKs, we sought to determine which RTKs are widely expressed in SCCOHT tumors. The expression of RTKs from RNA-Seq data on four SCCOHT primary tumors was hierarchically clustered to identify the top expressed RTKs in these cancers (Figure 3A). The top expressed genes across the tumors included two known ponatinib targets, *PDGFRA* and *FGFR1*, in addition to other RTKs such as *CSF1R, EPH5A, NTRK2*, and *ERBB3.* To functionally confirm that ponatinib inhibits signaling downstream of RTK activation, phosphorylation of p38 and Akt was examined in BIN67 cells (Figure 3B) which demonstrated a reduction in phosphorylation of p38, Akt S473 and Akt T308 following treatment with IC_50_ ponatinib (Figure 3B).

**Figure 3.**
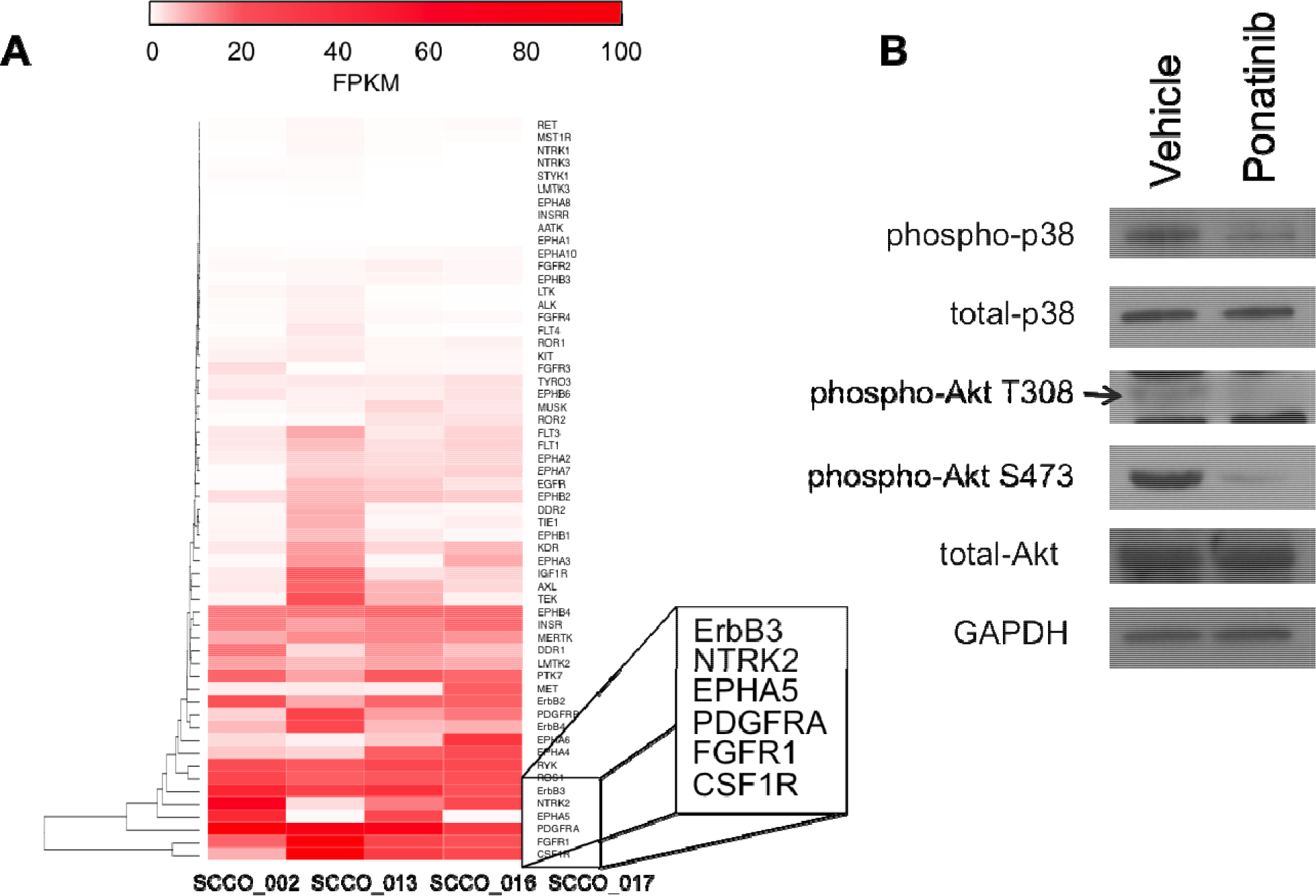
RTK expression in SCCOHT primary tumors 449 and downstream RTK signaling inhibited by ponatinib in SCCOHT cells. (A) Expression of all RTK genes in 4 SCCOHT primary tumors from RNA-Seq. Expression is displayed in FPKM, and genes are hierarchically clustered. The 6 most highly expressed RTKs are in inset. (B) Phosphorylation of signaling components downstream of RTKs are inhibited by ponatinib within 30 min treatment in BIN67 cells.

### Functional characterization of ponatinib action reveals broad kinase targeting in SCCOHT cells

Since ponatinib is a broad RTKi, several orthogonal approaches were undertaken to identify kinases inhibited by ponatinib in SCCOHT cells. First, RTK profiler arrays were employed to screen for top RTKs inhibited by ponatinib treatment in BIN67 and SCCOHT-1. This assay is a membrane-based sandwich immunoassay that captures key RTKs and detects differential pan-tyrosine phosphorylation in cell lysates across conditions. The quantitation of these arrays is shown in Figure 4A. Of the known ponatinib targets (highlighted in yellow), PDGFRα phosphorylation was both detected at a high level relative to other RTKs and also strongly inhibited by ponatinib in BIN67 cells only, but not in SCCOHT-1 cells. This discordance between BIN67 and SCCOHT-1 would suggest that ponatinib inhibits unidentified kinases in SCCOHT cells to affect cell growth. Among previously unreported targets of ponatinib, phosphorylation of EGFR in BIN67 and SCCOHT-1 cells was greatly inhibited by ponatinib, which represented the only common candidate between the two cell lines. Other RTKs strongly inhibited by ponatinib treatment in BIN67 cells alone include RYK, CSF1R, ALK, and TEK (Figure 4A).

**Figure 4.**
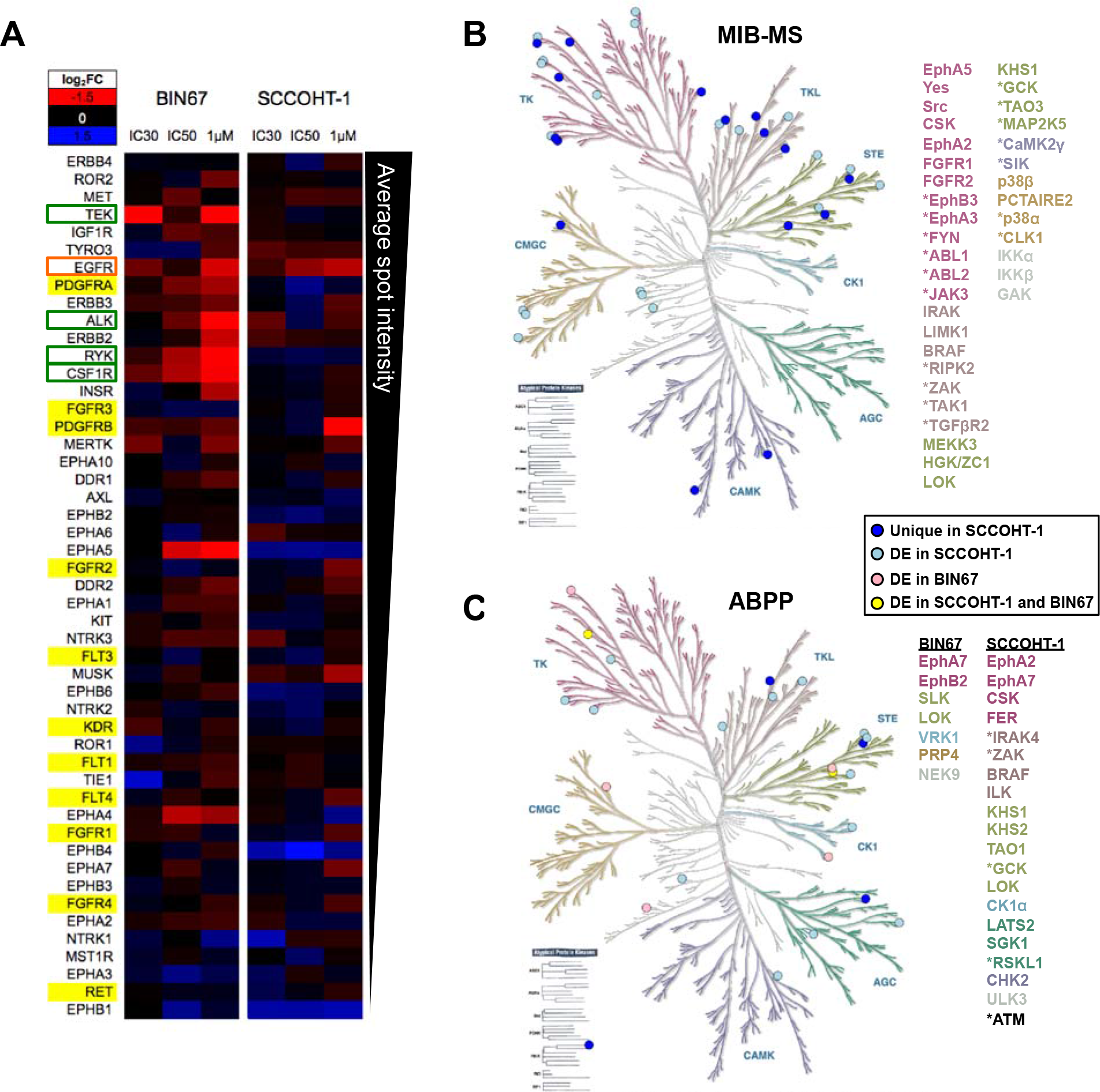
RTK signaling inhibited by ponatinib in SCCOHT cells. (A) Phospho-RTK array performed on lysates from BIN67 cells treated with ponatinib for 1 hour. Background normalized spot intensity relative to vehicle-treated control is represented as a heatmap, where blue indicates an increase in phosphorylation and red indicates a decrease. RTKs are arranged from top to bottom based on mean basal dot intensity. Known ponatinib targets are highlighted in yellow, kinases inhibited in both cell lines are outlined in orange and kinases inhibited in one cell line only are outlined in green. (B and C) Kinases identified from MIB-MS assay (B) and (C) ABPP assay in SCCOHT-1 cells (blue circles, 2μM, biological triplicates) and BIN67 cells (red circles, IC50, technical replicates) or in both cell lines (yellow circles) mapped to kinase dendrogram. The gene lists are color-coded based on corresponding kinase family in the dendrogram. Atypical protein kinases are listed in black. Differentially expressed kinases are shown in lighter shaded circles. Uniquely expressed kinases are shown in darker shaded circles and are marked with an asterisk in the gene list.

Given the divergent results of the RTK array assay between ponatinib sensitivity in SCCOHT-1 and BIN67, two additional proteomic approaches (MIB-MS and ABPP) were undertaken to more fully characterize functional inhibition of kinases by ponatinib. Multiplexed kinase inhibitor bead-mass spectrometry (MIB-MS) uses beads to pull down kinases from a cell lysate in the presence or absence of an inhibitor such as ponatinib (35,36). Binding of ponatinib to the kinase inhibits kinase binding to MIBs. Therefore, peptides corresponding to kinase targets are lost in the subsequent mass spectrometry analyses and thereby considered hits. Due to ponatinib hypersensitivity in SCCOHT-1 cells, this line was tested using MIB-MS to identify additional candidates for ponatinib targeting (full data in Supplemental Table 4). Kinase hits from most of the major kinase families were identified in SCCOHT-1 by this approach (Figure 4B). Kinases were considered targets in this experiment if, in the vehicle control, they were overexpressed (19 kinases) or uniquely detected (24 kinases) (Figure 4B). FGFR1, FGFR2, and Src are known ponatinib targets and were among the differentially expressed genes. These kinases similarly displayed reduced phosphorylation in the RTK array, but only at a high dose comparable to that used for the MIB-MS analysis, suggesting that other kinases might be more strongly inhibited by ponatinib at the relevant dose for cell death. Of the RTKs highly expressed in SCCOHT tumors (Figure 3A), EphA5 was also identified as a target of ponatinib in the MIB-MS experiment (Figure 4B).

A complementary approach to determination of those kinase upon which ponatinib operates in SCCOHT, Activity-Based Protein Profiling (ABPP), uses chemical probes that covalently bind to the ATP-binding pocket of kinases and therefore can tag kinases with accessible and conformationally active ATP-binding pocket in a protein lysate prior to mass spectrometry (full data in Supplemental Tables 5 and 6) (37). The same cutoff parameters described above for MIB-MS were used for this dataset to determine unique and differentially expressed active kinases. In SCCOHT-1 cells, we identified 13 differentially expressed kinases and 4 unique kinases in the vehicle control treated cells (Figure 4C). Only seven differentially expressed kinases in BIN67 cells were identified (Figure 4C). Two overlapping kinases were identified in both BIN67 and SCCOHT-1 cells: tyrosine kinase EphA7 and STE-family kinase LOK (Figure 4C). Interestingly, no RTK was significantly altered with ponatinib treatment in either cell line, which might represent a bias of this assay due to exclusion of active-site peptides via complementarity to other RTKs. Generally, tyrosine kinases and STE kinases were well-represented in both cell lines.

### Ponatinib is effective in animal models of SCCOHT

To determine ponatinib’s promise as a potential therapy, we tested its efficacy in animal xenograft models of SCCOHT. Mice bearing subcutaneous SCCOHT-1 xenograft tumors demonstrated an initial inhibition of tumor growth following ponatinib treatment, delaying tumor progression (Figure 5A). Tumor doubling times from 200 to 400mm^3^ in each group were 1.88±0.64 days in vehicle treated mice and 7.38±1.06 days in ponatinib treated mice (p-value < 0.0001). Treatment of SCCOHT-1 tumor bearing mice with ponatinib similarly improved median survival by 44% (Figure 5B). To further assess the efficacy of ponatinib in patient-derived tumor models, we used two SCCOHT patient-derived xenograft (PDX) models: PDX-465 and PDX-040. After 30 days of treatment, tumor growth rate was slowed in both PDX models, with statistically significant 58.6% and 42.5% decreases in tumor size relative to vehicle treatment in PDX-465 and -040, respectively (Figures 5C and 5D).

**Figure 5.**
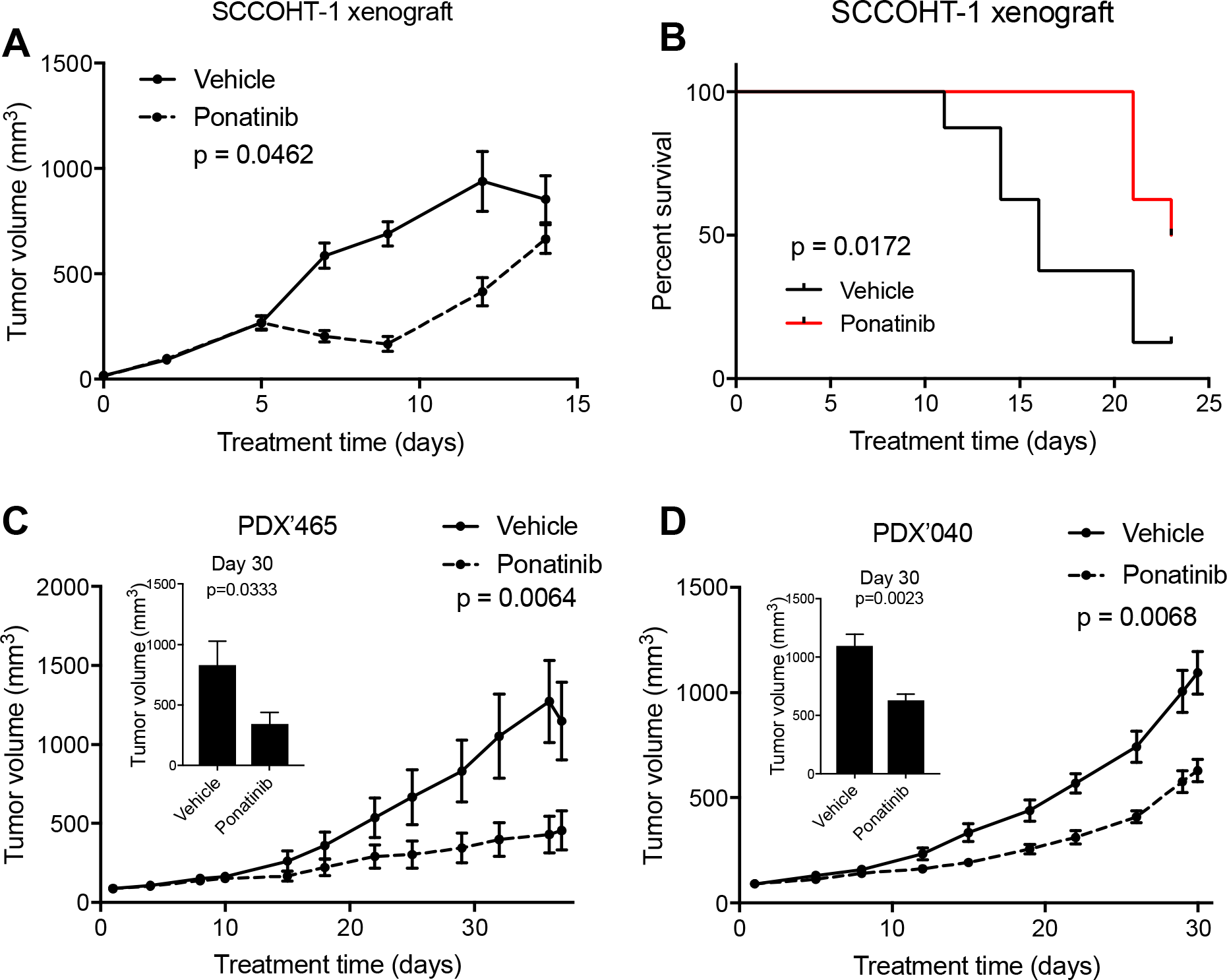
Efficacy of ponatinib in animal models of SCCOHT. (A) SCCOHT-1 xenograft growth with treatment with vehicle or ponatinib. (B) Survival of SCCOHT-1 xenograft-bearing mice following treatment with vehicle or ponatinib. (C and D) Tumor volumes of SCCOHT PDX model ‘465 (C) or ‘040 (D) growth with treatment with vehicle or ponatinib. Insets in (C) and (D) show tumor volumes at day 30.

### Discussion

High recurrence rates, extreme toxicity, and poor overall survival in SCCOHT patients treated with standard high-dose chemotherapy and radiation necessitate the identification of effective targeted therapies for this disease. Discovery of universal disruption of SWI/SNF complex function in SCCOHT through the loss of ATPase subunits SMARCA4 and SMARCA2 has made identification of targeted therapeutic vulnerabilities possible. Further, several novel approaches to selective killing of SWI/SNF-mutant cancers have recently been explored. The SWI/SNF complex normally functions to catalyze the movement and ejection of histones to regulate the accessibility of chromatin to gene expression machinery. Antagonism between SWI/SNF and the Polycomb Repressive Complex 2 (PRC2) described in *Drosophila* (38), has been recapitulated in human cells where it has been specifically shown that loss of SWI/SNF function leads to a dependence on the repressive functions of PRC2 (39). For this reason, inhibitors of the PRC2 catalytic subunit, EZH2, have been examined in SMARCB1-mutant rhabdoid tumors, ARID1A-mutant ovarian clear cell carcinomas, SMARCA4-mutant non-small cell lung cancers, and SMARCA4-mutant SCCOHTs (24,25,39-42). Such agents are now being examined in Phase I and II clinical trials in SMARCB1-negative or rhabdoid-like tumors, such as MRTs and SCCOHTs, but have not yet been approved for these indications. The SWI/SNF complex itself has also been identified as the major target of bromodomain inhibitors through direct interaction with bromodomain-containing SWI/SNF ATPases SMARCA2 and SMARCA4 (43). This is a particularly attractive target in some SMARCA4-deficient cancers in which residual SMARCA2 compensates in the absence of SMARCA4 as the functional SWI/SNF ATPase (44). Such an approach has been shown to be ineffective in SCCOHT, however, where expression of both SWI/SNF ATPases has been lost (3).

The effects of PRC2 dependence on tumor suppressor silencing in SWI/SNF-mutant cancers are well-studied. However, little is yet known about the effects of SWI/SNF dysfunction on the overexpression of oncogenes such as RTKs, which might cause a tumor to develop unique and targetable dependencies. Here, we show data supporting that targeting RTK activation in tumors driven by epigenetic defects may provide clinical benefit. Using HT siRNA and drug screens, our work has charted novel therapeutic vulnerabilities in SCCOHT cells. These studies have not only confirmed *in vitro* sensitivity to cytotoxic drug classes already utilized in SCCOHT treatment (e.g. tubulin inhibitors and DNA damaging agents), but they have also pinpointed novel drug classes, such as cardiac glycosides, not previously implicated in treatment of SWI/SNF-mutant cancers. Overall, these data converge on identification of hypersensitivity to the FDA-approved multi-kinase inhibitor ponatinib and thereby emphasize the importance of RTK signaling in these epigenetically dysregulated tumors.

We generally observed a striking similarity in drug sensitivity across the SCCOHT cell lines and SVOG3e. While this observation strengthens the case for the sensitivity of SCCOHT tumors to these agents, the similar sensitivity of SWI/SNF-intact SVOG3e cells indicates that most of these agents do not selectively target SWI/SNF-mutant cells. The cell of origin of SCCOHT remains unclear, but current understanding suggests derivation from a germ cell lineage (45) distinct from the cells from which SVOG3e cells were derived. Further work must expand cell models for each agent to better understand the relationship between cell lineage and drug response. Interestingly, previous reports have shown expression of PDGFRs and c-kit, known ponatinib targets, in granulosa cell tumors (46), suggesting convergent singaling dependencies that may account for sensitivity to this drug in SCCOHT and SVOG3e cell lines. Additional work is required to more completely characterize vulnerability of cancers with SWI/SNF loss to the agents identified in this screen.

In examining overlap between the 42 hits in the drug screen and the 109 hits in the siRNA screen, RTKs were the primary point of intersection. In addition to the overlap seen within our HT screens, previous work has demonstrated sensitivity of SWI/SNF-mutant MRTs to ponatinib (30). Mechanistically, our work supports data reported in SMARCB1-mutant MRTs, in which loss of the core SWI/SNF subunit SMARCB1 results in elevated FGFR or PDGFR expression (26,30). We similarly observed sensitivity of SCCOHT cell lines to ponatinib, a broad RTKi. Multiple methods were integrated in an attempt to functionally identify a common target of ponatinib responsible for efficacy of this drug in SCCOHT. Criteria considered included: 1) expression in SCCOHT primary tumors, 2) requirement of expression for cell growth, and 3) functional inhibition in the presence of ponatinib. Using these criteria, however, no common target was identified, both between the extensive methods described here and the two SCCOHT cell lines. Furthermore, the sensitivity of SCCOHT cells to other RTKi is inversely related to the specificity of the RTKi to its intended target. Together, this evidence suggests general reliance of SCCOHT on multiple kinases such that a broad kinase inhibitor such as ponatinib is effective.

When taken in combination with similar findings in MRTs (26,27,30) and the sensitivity observed in other SWI/SNF mutant lung cancer cell lines reported here, we believe drug sensitivity findings in the SCCOHT and MRT models might be more broadly applicable to a larger set of “SWI/SNF-omas”, or cancers with mutations in the SWI/SNF complex. This group accounts for up to 20% of all cancers, and therefore has potential for broader translational relevance. Our work also demonstrates the efficacy of ponatinib in SCCOHT preclinical animal models. Ponatinib is a clinically approved agent used in the treatment of leukemias, and is being assessed for efficacy in a number of other cancers, including solid tumors, in ongoing clinical trials. This work demonstrates promise for ponatinib in improving outcome for SCCOHT patients as well.

## Acknowledgements

This work is supported by the National Institutes of Health (1R01CA195670-01, to D.G.H., J.T. and B.W.), the Canadian Cancer Society Research Institute (#703458, to D.G.H.), the Terry Fox Research Institute Initiative New Frontiers Program in Cancer (D.G.H.), the Marsha Rivkin Center for Ovarian Cancer Research, the Ovarian Cancer Alliance of Arizona, the Small Cell Ovarian Cancer Foundation, and philanthropic support to the TGen Foundation. We would also like to thank the SCCOHT patients, their families and communities, and the clinicians who have contributed significantly to the motivation and feasibility of this work.

